# High-resolution mapping of foveal vision

**DOI:** 10.1101/2025.10.08.681250

**Authors:** Lynn Schmittwilken, Michele Rucci

**Affiliations:** Brain and Cognitive Science, University of Rochester, New York; Computational Psychology, Technische Universität Berlin, Germany

## Abstract

Human vision critically relies on the foveola, a tiny region of the retina with the highest photoreceptors density. Although it spans only *∼*0.1% of the visual field, the foveola accounts for nearly one third of projections to the visual cortex. Accurate assessment of foveal function is therefore essential for monitoring visual processes and visual health. However, probing this region is challenging because of its minute size and the incessant eye movements that humans perform, even when fixating on a single point. Here, we leverage recent advances in eye-tracking and gaze-contingent display control to develop a visual field test capable of mapping foveal sensitivity with high precision and reliability. Applied to healthy observers, this method reveals substantial idiosyncratic variability both across and within individuals, with peak sensitivity consistently shifted toward the temporal visual field. By enabling high-resolution individualized assessment of foveal vision, this approach opens new avenues for both clinical diagnostics and basic research.

## 1 Introduction

The foveola, located at the very center of the retina, is critical for human visual processing (Rentschler & Treutwein, 1985; Toet & Levi, 1992; Poletti, 2023, for review) and a wide range of visual behaviors such as reading and driving (Legge, Rubin, Pelli, & Schleske, 1985; Daien et al., 2014). Although it spans only 1° of visual angle—the extent of the index fingernail at arm’s length—the foveola maps onto a major portion of the visual cortex (Daniel & Whitteridge, 1961; Tootell, Silverman, Switkes, & De Valois, 1982), and acuity declines rapidly with distance from the foveola (Fig. 1A-B; Weymouth, 1958; Green, 1970).

**Figure 1:**
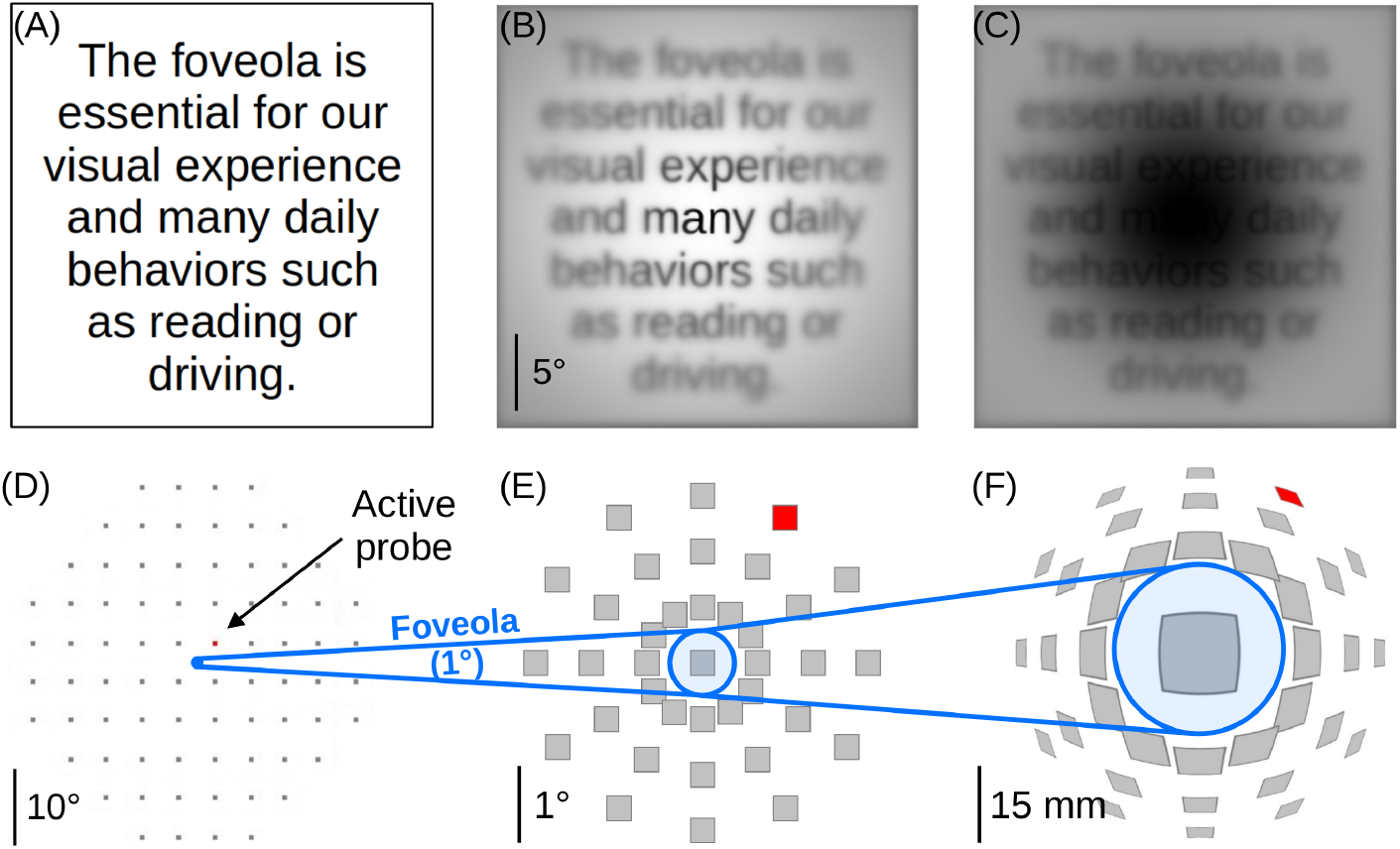
Despite its relevance, the foveola is not represented in standard visual field tests. An example text image (*A*) and how it appears (*B*) when sampled with retinal resolution, which declines away from the foveola. (*C*) Macular degeneration, a condition marked by central vision loss, illustrates the profound impact of impaired foveal vision. (*D*) Standard visual field (perimetry) methods assess sensitivity too coarsely to map foveal functions. These tests sequentially probe sensitivity my presenting brief flashes at various locations. (*E*) Even in tests that use high density probe grids, like the macular integrity assessment microperimeter, the foveola is represented by only a single data point. (*F*) Yet, the foveola is disproportionally represented in the visual cortex. The map shows how much the central portion of *E* is distorted by cortical magnification. The red probe marks a location 3° away from the center.

Given the foveola’s pivotal role in visual processing, monitoring its function is essential in both scientific and clinical contexts (cf. Qian et al., 2024). A relevant example that demonstrates the importance of monitoring foveal vision is macular pathology, the leading cause of blindness and low vision in the Western world (Hazel, Petre, Armstrong, Benson, & Frost, 2000). Macular pathologies are characterized by a progressive central vision loss (Fig. 1C). Early detection is critical to prevent irreversible damage, yet structural changes often become visible only after they extended well beyond the foveola (Sunness et al., 1999; Cassels, Wild, Margrain, Chong, & Acton, 2018).

Emerging evidence suggests that subtle changes in foveal vision can already arise before structural abnormalities become detectable (Vullings & Verghese, 2021; Clark, Moon, Jenks, Kapisthalam, & Poletti, 2023). This offers a valuable window for early intervention. However, standard tools to assess visual function, such as visual field tests (*visual perimetry*), are not yet suited to detect such early changes as they often occur at a too fine spatial scale on the order of a few arcminutes 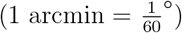. Standard perimetry typically samples large portions of the visual field (up to 60° of visual angle) with a coarse spatial resolution (Fig. 1D; Johnson, Wall, & Thompson, 2011; Phu et al., 2017). Even perimetry systems that target the central visual field typically group the entire foveola into a single measurement point (Fig. 1E; Charng et al., 2020). This makes existing perimeters too coarse to detect early sensitivity changes within the foveola, despite its disproportionate relevance for visual processing (Fig. 1F).

Probing foveal vision is challenging due to the foveola’s minute size and the constant presence of involuntary eye movements. Even during attempted fixation, the human eye is never still (Fig. 2). Humans perform rapid microsaccades several times per second, interspersed with slower, erratic drifts (Ratliff & Riggs, 1950; Rucci & Victor, 2015). These movements introduce uncertainty about the stimulus’ position on the retina (Fig. 2B-C; Cherici, Kuang, Poletti, & Rucci, 2012), and prevent detecting subtle changes in foveal sensitivity during natural vision (Poletti, Listorti, & Rucci, 2013; Clark et al., 2023). As a result, sensitivity changes in the foveola go undetected unless eye movements are actively accounted for during testing. Most eye trackers, however, suffer from limited resolution and accumulating localization errors over time (Kimmel, Mammo, & Newsome, 2012; Bowers, Gautier, Lin, & Roorda, 2021; Carr, Pescuma, Furlan, Ktori, & Crepaldi, 2022) that severely hinder assessment of foveal function.

**Figure 2:**
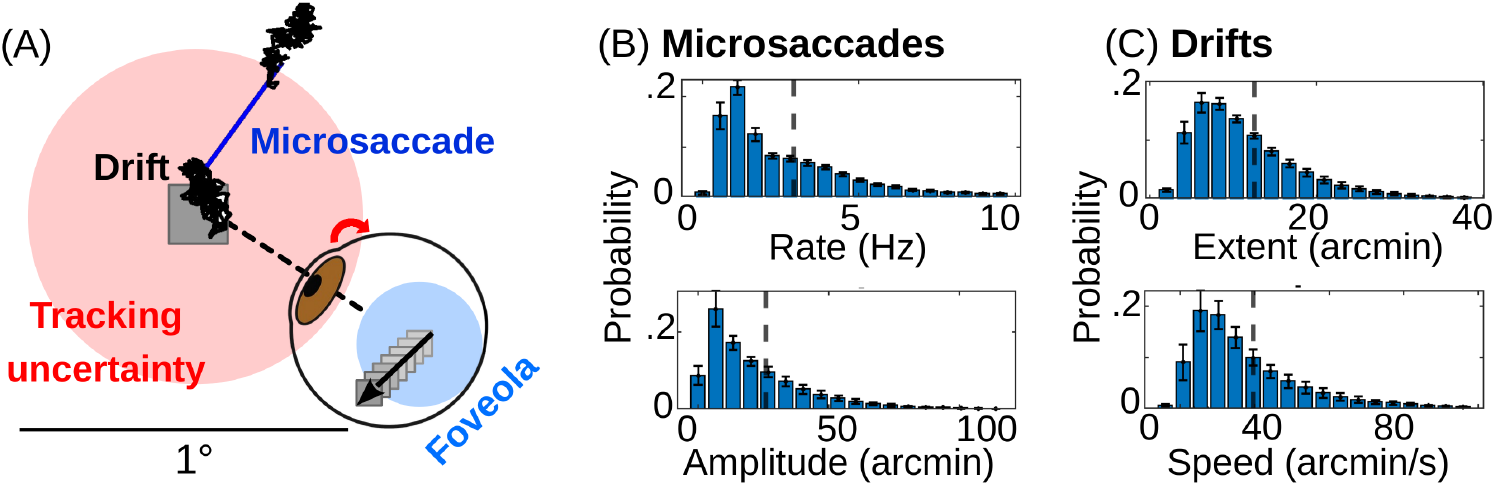
Eye movements challenge mapping the foveola. (A) Eye movements continually occur during fixations and can shift a fixated probe presented to the center of the foveola fully outside of it. These movements occur at the lower limit of what standard eye trackers can trace. They are divided into fast microsaccades (blue) and slower drifts (black). (B-C) Eye movement statistics during fixations, such as in a standard perimetry test. Observers perform 2-3 microsaccades per second that shift their foveola by 23.5+-19.7 arcmin. In addition, drifts shift the foveola by 12.4+-6.5 arcmin per fixation with a velocity of 39.4+-14.7 arcmin/s. Histograms show mean probabilities and standard deviations across N=11 observers.

Here, we describe a perimetry method that builds on recent advances in eye-tracking hardware, custom calibration procedures, and gaze-contingent stimulus control to precisely probe selected retinal locations during normal fixational eye movements. By means of rigorous psychophysical testing, we show that this method enables reliable, fine-grained mapping of foveal sensitivity.

## 2 Results

### 2.1 A gaze-contingent approach

**Fast and precise eye tracking.** Most standard perimetry systems monitor fixational stability with corneal-reflection-based eye trackers. These systems, which typically estimate gaze from the displacement between the first Purkinje image of an infrared beam and the pupil, have relatively coarse spatial (1–2°) and temporal (30–100 Hz) resolution (Heijl, Patella, & Bengtsson, 2021). Even newer microperimeters that use fundus-image tracking, i.e. continuously image the retina by means of a scanning laser ophthalmoscope, operate at too coarse spatial (0.25–1°) and temporal (25–50 Hz) resolutions (Pfau et al., 2021; Yang & Dunbar, 2021). These constraints make it difficult to reliably resolve visual sensitivity within the foveola in the presence of incessant eye movements (Fig. 2).

To overcome these limitations, we used a digital dual-Purkinje image (dDPI) eye tracker (Fig. 3A), which estimates eye position by measuring the relative displacement of reflections from multiple surfaces of the eye: the cornea and the posterior lens (Wu et al., 2023). This technique has long been regarded as a gold standard in oculomotor research for its precision and robustness (Cornsweet & Crane, 1973; Crane & Steele, 1985). Until recently, traditional DPI systems required complex analog setups, and were therefore limited to specialized laboratories (Deubel & Bridgeman, 1995; Hawken & Gegenfurtner, 2001). However, recent advances have led to the development of a digital system, which preserves its precision with improved usability (Wu et al., 2023). Unlike most commercial eye trackers, the dDPI system is fully transparent, as users have access to the raw, unfiltered data without proprietary smoothing or filtering. We validated its subarcminute precision and high temporal resolution (1 kHz) with an artificial eye on a motorized stage (Fig. 3B).

**Figure 3:**
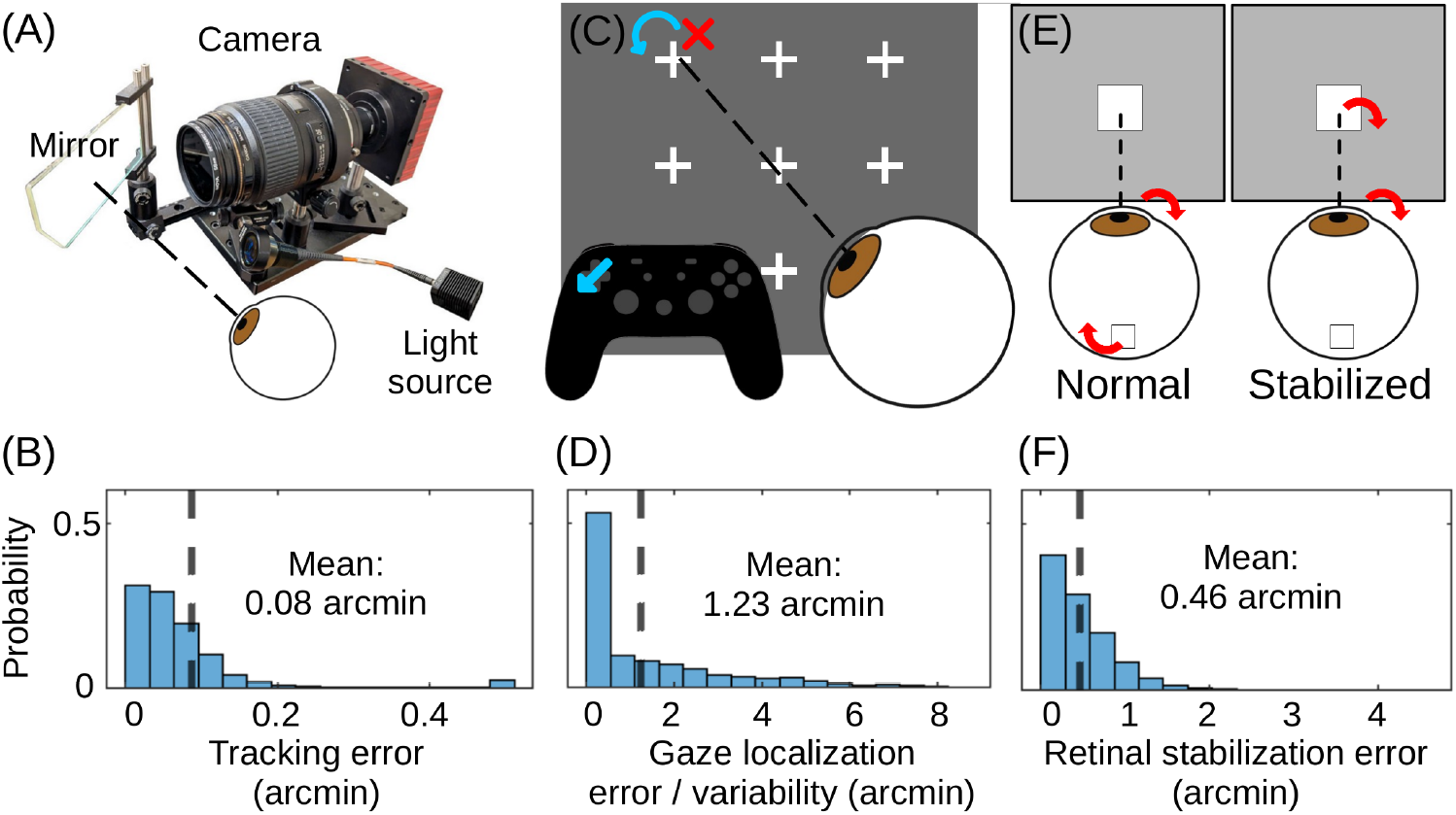
High-precision integration of digital dual-Purkinje eye tracking, gaze calibration, and real-time compensation of eye movements. (*B*) Accuracy of the dDPI system (*A*) measured with an artificial eye that was mounted on a rotational stage. The histogram shows the deviation between the physical and estimated displacements. (*C*) Gaze calibration was performed in two steps: subjects first fixated a grid of nine equidistant points, allowing estimation of gaze on the display. They then refined these estimates (red cross) until they matched their perceived center of gaze using a controller. (*D*) Residual offsets between measured gaze and the subject’s perceived center of gaze during repeated fixations. (*E*) To deliver probes to targeted retinal locations despite natural eye movements, we implemented real-time gaze-contingent stimulus control. (*F*) Difference between actual and compensated gaze position, which results from the delay between gaze sampling and display update.

We have previously shown that the dDPI reliably captures fine-scale eye movements across observers (Intoy, Mostofi, & Rucci, 2021; Clark, Intoy, Rucci, & Poletti, 2022). These capabilities provide the precision necessary to track gaze with the spatial and temporal fidelity required for mapping visual sensitivity across the foveola.

**Accurately mapping gaze relative to the preferred retinal locus.** Precise mapping of foveal vision requires knowing not just how the eyes move, but where the observer is looking on the display at any given moment. This step requires to translate eye tracker signals into spatial coordinates on the display, which is a key limiting factor for high-precision eye tracking. Even though some modern video eye trackers might in theory be able to measure gaze displacements with high precision (*∼*1 arcmin), their built-in calibration procedures often yield spatial inaccuracies of up to 1° (SR Research, 2005, *2013; VPixx Technologies, 2020)*, which is too coarse to probe sensitivity within the foveola.

To overcome this limitation, we implemented a custom dual-step calibration procedure that achieves arcminute accuracy and aligns gaze localization with the observer’s perceived center of gaze. In the first calibration phase, participants sequentially fixate on nine probes arranged in a uniform grid (Fig. 3C). This data is then used to generate an initial estimate of the gaze-to-display mapping. In the second phase, subjects refine these estimates with a controller to adjust the position of a visible marker (a red cross) to match the location they perceive as the exact center of their gaze.

This refinement serves two purposes. First, it improves localization accuracy by an order of magnitude, yielding a mean error of just 1.23 arcminutes (0.02^°^; Fig. 3D), consistent with previous work (Poletti et al., 2013). Second, it anchors the coordinate system to the perceived center of gaze, also known as the preferred retinal locus of fixation, instead of relying on anatomical or optical landmarks like the anatomical center or peak cone density (Wang, Cherici, & Rucci, 2025). These landmarks are often not easily accessible in behavioral experiments and do not match where observers feel they are looking (Putnam et al., 2005; Kilpeläinen, Putnam, Ratnam, & Roorda, 2021).

This distinction is especially important for our high-resolution perimetry. Using the same perceptually defined origin across participants likely reduces variability introduced by differences in fixation strategy and oculomotor behavior. Our method ensures that visual sensitivity is mapped relative to the retinal location that the observer naturally uses to explore the world, providing both functional relevance and measurement consistency across sessions and individuals.

**Compensating eye movements in real-time.** To measure foveal sensitivity with high precision, it is desirable to stimulate specific retinal sites in a controlled and repeatable man-ner. However, even the smallest fixational eye movements cause stimuli to shift unpredictably across the foveola (Fig.2B–C). Without real-time compensation (Fig. 3E), it therefore becomes difficult to ensure that a stimulus consistently engages the intended retinal site.

To address this challenge, we incorporated EyeRIS, an established real-time gaze-contingent display system that compensates for eye movements as they occur (Santini, Redner, Iovin, & Rucci, 2007). In our setup, EyeRIS runs on a real-time Linux system with custom I/O boards and shares the same host computer and GPU as the digital dual-Purkinje image (dDPI) eye tracker (Wu et al., 2023). This tight integration ensures fast and consistent data transfer, with end-to-end delays of less than 5 ms (two frames at 360 Hz). As a result, EyeRIS updates stimulus positions on the display based on the most recent eye position with a spatial precision of 0.46 arcminutes (=0.008°; Fig.3F). This accuracy ensures that probes remain stably aligned with the intended foveal location, even in the presence of incessant eye movements.

Together, the precision of our eye tracking, gaze localization, and real-time stabilization allows us to target foveal locations with a spatial accuracy of better than 2 arcminutes. This resolution approaches the minimum angle of resolution (0.5 arcmin; Rossi & Roorda, 2010) given the photoreceptor density in the foveola. Such fine-grained control enables reliable stimulation of specific retinal sites, even in the presence of continuous fixational eye movements.

### 2.2 High-resolution mapping of sensitivity across the foveola

Leveraging the high precision of our system, we mapped visual sensitivity at 13 discrete retinal locations within the central 1° of the visual field (Fig. 4A). Probe size was 5 arcmin, and probe sites were spaced 7 arcminutes apart, allowing dense sampling across the foveola.

**Figure 4:**
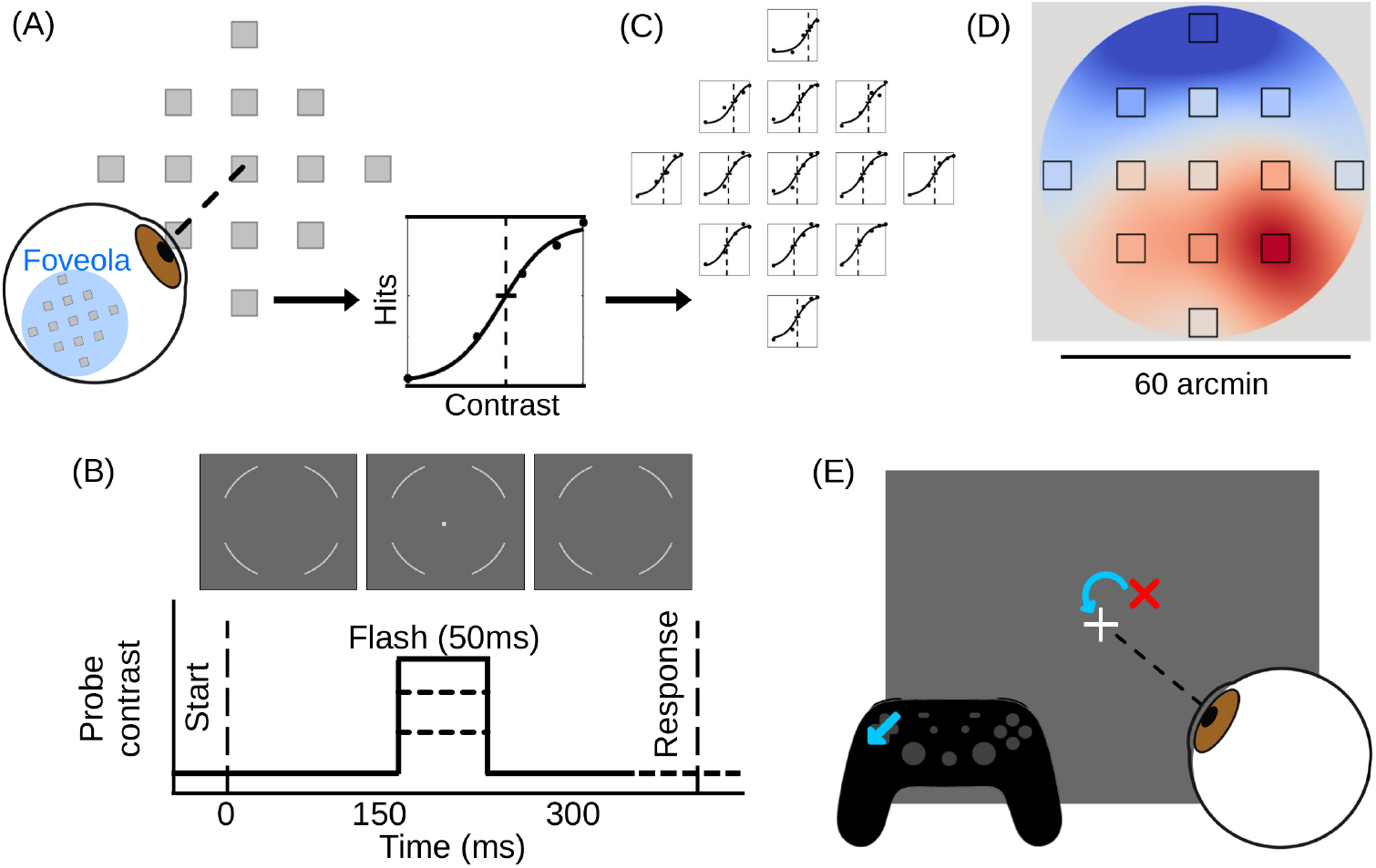
Mapping foveal sensitivity. (A) Sensitivity was measured at 13 retinal sites within 60 arcminutes of subjects’ perceived center of gaze. Probes (5 arcmin) were spaced 7 arcmin apart to densely sample the foveola. (B) On each trial, a probe appeared with 50% probability, 150 ms after trial onset, at one of the predefined locations. Probe contrast (Weber contrast: ratio of probe to background luminance) varied across trials. Probes were gaze-contingently stabilized to ensure precise retinal stimulation. (C) Psychometric functions were fit to detection responses at each location to extract contrast thresholds (50% hit rates). (D) Thresholds were interpolated using cubic splines to generate continuous sensitivity maps across the foveola. (E) Brief recalibration trials were interleaved every 15 trials to maintain alignment between measured gaze position and subjects’ perceived center of gaze.

On each trial, subjects fixated on a uniform gray background (10 cd/m^2^), aided by peripheral white arcs at 2.5° eccentricity that facilitated stable fixation without interfering with foveal vision (Fig. 4B). Trials began with a button press and an auditory cue which signaled probe onset. After 150 ms, a probe was presented for 50 ms at one of the 13 predefined retinal locations with a 50% probability of appearance. Within 13 trials, probes were presented at each site only once. Subjects indicated whether the probe was present or absent and received auditory trial-by-trial feedback.

Contrast sensitivity was quantified by fitting psychometric functions to the detection rates at each retinal location (Fig. 4C). We extracted contrast thresholds corresponding to 50% hit rates and applied cubic interpolation across retinal locations to generate continuous maps of foveal sensitivity (Fig. 4D). A mild Gaussian filter was applied to reduce spatial discontinuities and reveal fine-grained variations in sensitivity within this highly specialized retinal region.

To prevent cumulative errors in gaze localization over the course of testing (Kimmel et al., 2012; Bowers et al., 2021; Carr et al., 2022), we interleaved brief recalibration trials every 15 trials. During these, subjects viewed their estimated gaze position and adjusted it to align with a central fixation marker (Fig. 4E), following the same procedure as during initial calibration (Fig. 3C). These periodic adjustments were used to ensure continued alignment with each subject’s perceived center of gaze and maintained the high tracking accuracy necessary for precise stimulus delivery.

### 2.3 Idiosyncratic maps of visual sensitivity

To test the efficacy of our method, we mapped visual sensitivity across the foveola of 11 healthy subjects. Thresholds varied markedly across individuals (Fig. 5A), with the most sensitive participants outperforming the least sensitive by a wide margin. These interindividual differences highlight the utility of our approach for capturing fine-grained, subject-specific profiles of foveal function, even in healthy populations.

**Figure 5:**
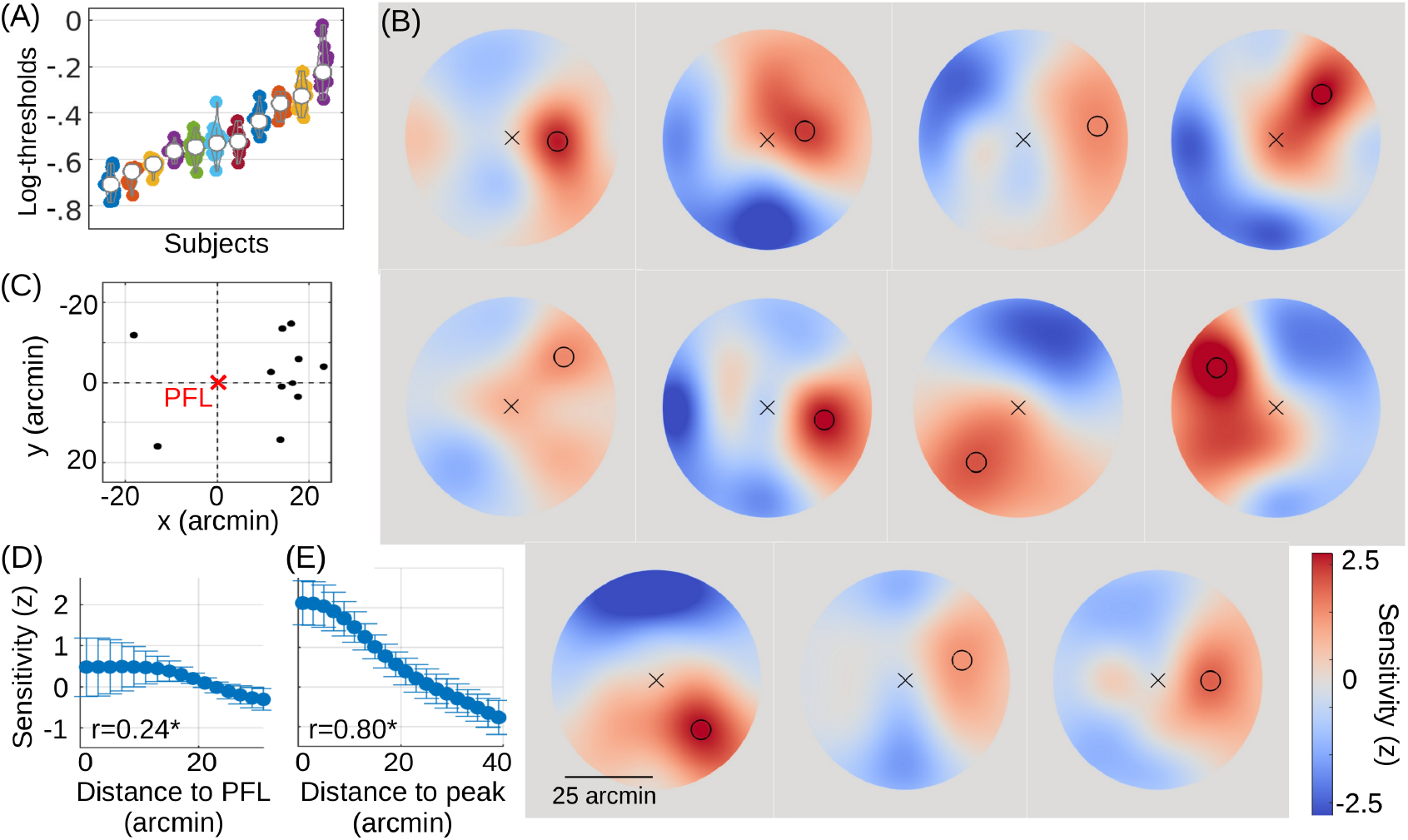
Sensitivity maps vary widely across individuals and space. (A) Contrast thresholds measured at the 13 foveal sites for each subject show substantial spatial and inter-individual variability. (B) Sensitivity maps for all *N* =11 subjects. Crosses indicate the preferred fixational locus (PFL); circles mark the retinal location with peak sensitivity. (C) Peak sensitivity locations are consistently offset toward the right (temporal) visual field in most observers. (D–E) Sensitivity declines systematically with distance from the PFL (D) and even more strongly with distance from the individual peak sensitivity location (E). Data show mean ± standard deviation across subjects, binned in 2 arcmin intervals. Pearson’s *r* indicates the correlation between sensitivity and distance.

To compare how sensitivity patterns varied across subjects, we computed z-scored threshold values. This normalization removes differences in overall sensitivity and yields a direct estimate of the reliability and effect sizes of our method. The resulting z-scored sensitivity maps (Fig. 5B) reveal highly individualized and non-uniform sensitivity profiles across the foveola. On average, thresholds at the most sensitive retinal location were 18% smaller than at the center of gaze, and 51% smaller than at the least responsive site. These variations challenge the long-held assumption that foveal sensitivity is spatially homogeneous and demonstrate that even within the foveola, fine-scale structure strongly shapes visual function.

In all subjects, the location of peak sensitivity was consistently offset from the preferred fixational locus, typically toward the temporal visual field (Fig. 5C). This directional bias suggests a systematic, retinotopically organized asymmetry within the central foveola.

Sensitivity declined reliably with distance from the preferred fixational locus (Fig. 5D; *r* = *−*0.28, *p* = 0.009, *R*^2^ = 0.076), and even more steeply with distance from the site of peak sensitivity (Fig. 5E; *r* = *−*0.80, *p <* 10^*−*5^, *R*^2^ = 0.643). On average, thresholds increased (i.e. sensitivity decreased) by approximately 35% per degree of visual angle from the site of peak sensitivity. These spatial gradients demonstrate that the foveola contains a finely structured sensitivity landscape, with measurable functional consequences at an arcminute resolution.

### 2.4 Assessing reliability

To evaluate the robustness of our method, we collected extensive data across multiple days from each subject and performed a series of analyses. Our approach enhances reliability through three key components: (1) gaze-contingent stimulus presentation, (2) continuous recalibration to maintain precise gaze localization, and (3) exclusion of trials containing microsaccades. We first assessed the individual contribution of each element.

As expected, eye position varied substantially during the measurement, spanning over 60 arcmin, i.e. the size of the entire foveola (Fig. 6A). Subjects recalibrated their gaze center approximately three times per minute, with shifts up to 20.25 arcmin between recalibrations (Fig. 6B). Simulations showed that omitting eye movement compensation and repeated recalibrations would underestimate foveal inhomogeneities by at least 58% ± 12% (relative difference between the absolute maxima of the two approaches; Fig. 6C).

**Figure 6:**
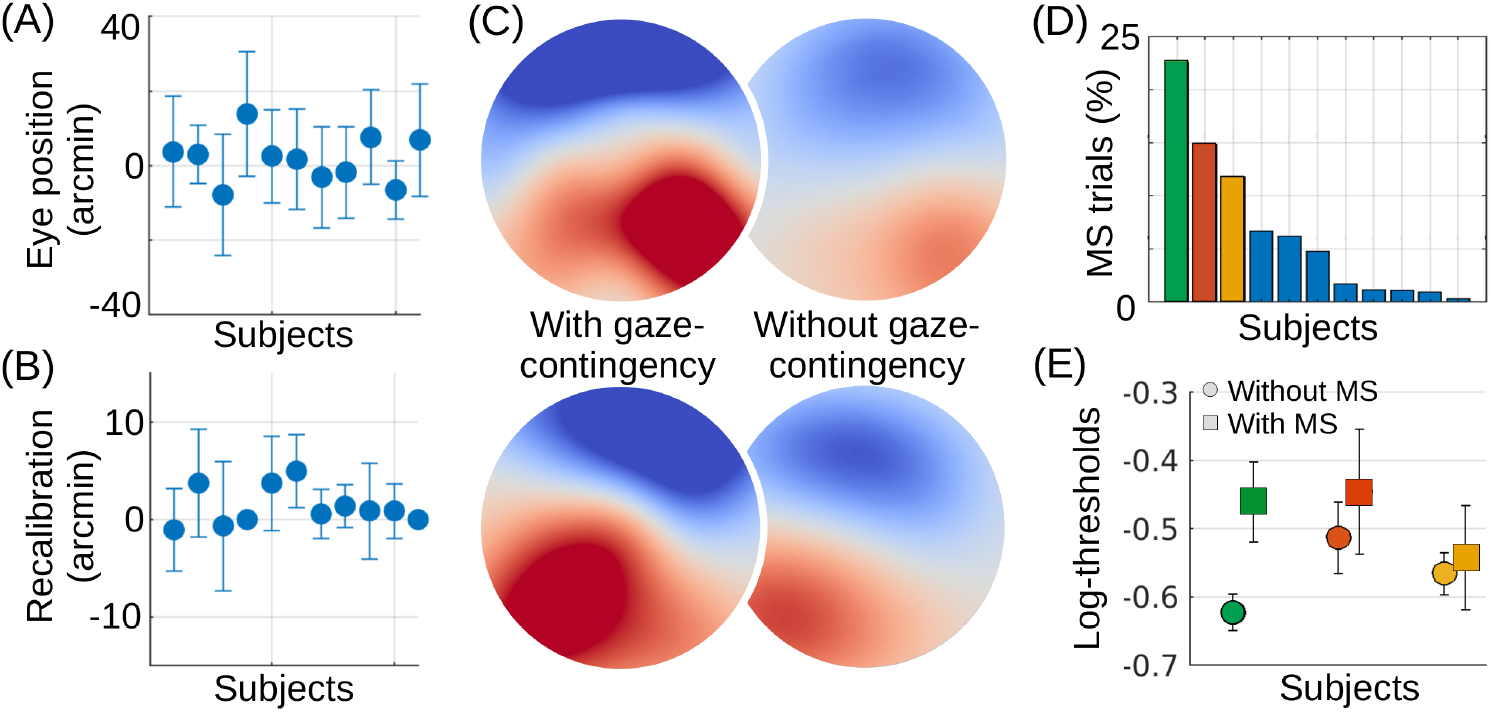
Importance of gaze-contingency and recalibration. (A) Eye position (mean±std) on the display for each subject, illustrating the extent of eye movements during attempted fixation. (B) Recalibration adjustments (mean±std) applied to maintain high-precision eye tracking across subjects. Both eye movements and recalibrations are substantial, emphasizing the necessity of compensating for them to reliably map foveal sensitivity. (C) Sensitivity maps from two representative subjects, showing the effect of omitting eye movement compensation. Without compensation, sensitivity maps appear 58±12% more homogeneous, calculated as the relative difference between maximum sensitivity of the original and uncorrected maps across all subjects. (D) Percentage of trials which contain at least one microsaccade during probe flashes for each subject. Some participants exhibited microsaccades in up to 23% of trials. (E) Perceptual thresholds estimated separately for trials with and without microsaccades in three subjects (green, red, and yellow) with sufficient microsaccade trials. Thresholds are notably higher and more variable in the presence of microsaccades, underscoring the importance of excluding these trials from analysis.

We removed microsaccades trials prior to computing foveal sensitivity. Microsaccade rates varied widely between subjects, with some exhibiting microsaccades during up to 23% of trials near probe presentation (Fig. 6D). Interestingly, these microsaccade rates were did not change much across sessions and were independent of observers’ expertise. The presence of microsaccades significantly increased threshold estimates and variability (Fig. 6E), underscoring the necessity of excluding these trials for accurate sensitivity mapping.

Next, we assessed the consistency of sensitivity maps across days. Sensitivity maps showed stable patterns over time (Fig. 7A for two representative subjects), indicating both the robustness of our method and the temporal stability of foveal sensitivity.

**Figure 7:**
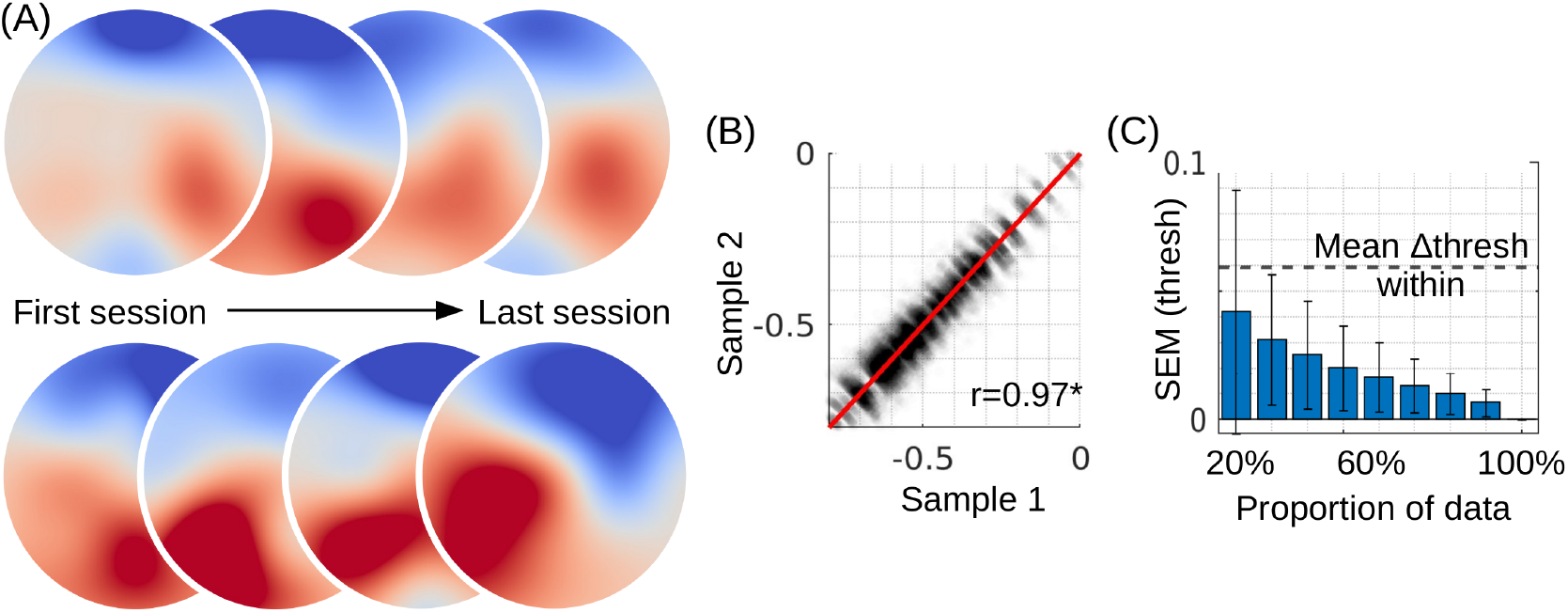
Reliability of the method. (A) Sensitivity maps are consistent across sessions, shown for two representative subjects. (B) For split-half reliability, we randomly split the datasets of each subject into two halves, and derived thresholds. We plot all thresholds derived from the first half (x-axis) against the thresholds from the second half (y-axis). Perfect alignment is indicated by the red diagonal. (C) Threshold SEMs are smaller than the threshold differences between probes *within* subjects (differences across subjects are much larger). This shows that we can reliably detect intraindividual differences with only 30% of the data.

To quantify this consistency, we performed split-half reliability analyses. Split-half reliability is a method to assess test reliability by dividing the empirical data into two halves and correlating the scores from each half to determine their similarity. In our case, we randomly split the datasets of each subject into two halves, and correlated the sensitivity estimates derived from the two halves (Fig. 7B). Split-half reliability revealed an excellent correlation (r=0.97), indicating that the method is highly reliable.

To test whether the sensitivity differences across space and subjects are statistically meaningful, we performed bootstrap analyses. These revealed that our method could reliably detect threshold differences as small as 0.015 Weber contrast across the foveola with as little as 30% of the data (Fig. 7C).

## 3 Discussion

We introduce a method to map visual function in the foveola, a region crucial for perception but historically underexplored due to technical limitations. As we demonstrate, the proposed approach is able to measure foveal sensitivity with unprecedented resolution and accuracy, integrating advanced display and eye tracking technologies with precise psychophysical tools. This innovation opens new avenues for both clinical and scientific investigations into central vision.

A major challenge for studying visual function inside the foveola is the continuous presence of eye movements, which occur even when observers fixate on a single spot (Ratliff & Riggs, 1950; Rucci & Victor, 2015). These eye movements shift fixated targets completely out of the foveola multiple times per second (Fig. 2). Although recent advances have been made in video-based eye tracking, the spatial resolution required to accurately track these movements remains elusive (Kimmel et al., 2012; Choe, Blake, & Lee, 2016; Bowers et al., 2021). In comparison, our system improves gaze-localization accuracy by an order of magnitude (Fig. 3), integrating three core innovations: (1) a custom-built dual-Purkinje image eye tracker that captures eye movements at 1 kHz with sub-arcminute precision (Fig. 3A, Wu et al., 2023), (2) repeated calibration procedures, which allow for gaze localization with arcminute accuracy (Fig. 3B-C), and automatically align eye traces to observers’ preferred fixational locus, and (3) real-time compensation of eye movements, enabling accurate stimulation of targeted foveal locations even while the eyes move (Fig. 3D).

Each component of our system was carefully chosen for its contribution to the overall performance and accuracy of the method. Importantly, we designed the system to be highly modular, creating a flexible platform that can incorporate technological advancements, particularly in eye tracking and psychophysical protocols, to enhance accuracy and efficiency as new innovations emerge.

To perform a perimetry within the foveola, it is mandatory to localize gaze with arcminute precision. We selected a dual-Purkinje image eye tracker because it provides the required precision (Wu et al., 2023) while preserving natural viewing conditions and being both user-friendly and scalable. In comparison, most other modern video-based eye trackers lack the requisite spatial precision (Kimmel et al., 2012; Bowers et al., 2021) and suffer from pupil artifacts (Choe et al., 2016). In contrast, eye tracking systems with a scanning laser ophthalmoscope could provide a real alternative to using a dual-Purkinje eye tracker (Stevenson, Roorda, & Kumar, 2010; Sheehy et al., 2012). From our point of view, however, video-based eye trackers currently still seem more accessible and less invasive.

For the psychophysical assessment, we used a constant stimuli approach to rigorously evaluate foveal sensitivity (Blackwell, 1952). Although adaptive psychophysical methods, such as SITA, have been successful in other perimetry applications (Bengtsson, Olsson, Heijl, & Rootzén, 1997; Bengtsson & Heijl, 1998), we chose method of constant stimuli to establish a reliable baseline first, as using faster methods usually has the drawback of reducing accuracy and reproducibility (Shirato, Inoue, Fukushima, & Suzuki, 1999). This choice provides the basis for a robust comparisons with other methods in the future. A long-term perspective for clinical applications could still be to incorporate adaptive methods to further enhance the efficiency of the proposed approach.

The role of eye movements in vision research has gained significant interest in recent years, with particular attention to their clinical implications (Lebedev, Belokopytov, Rozhkova, Vasilyeva, & Gracheva, 2024). The existing approach that most closely resembles our method, both in function and application, is *microperimetry*. Unlike standard perimetry, where eye tracking is either absent or too imprecise to account for small fixational shifts, microperimetry represents a major improvement by introducing real-time eye tracking and gaze-contingent stimulus presentation (Rohrschneider, Bültmann, & Springer, 2008; Acton & Greenstein, 2013). This allows stimuli to be accurately aligned with the intended retinal location, even in the presence of natural eye movements. These features allow visual sensitivity to be assessed relative to the current point of fixation, significantly improving spatial precision and test reliability (Laishram, Srikanth, Rajalakshmi, Nagarajan, & Ezhumalai, 2017; Pfau et al., 2021; Yang & Dunbar, 2021). While microperimetry systems also capture fundus images to correlate functional sensitivity with retinal structure, it is the integration of gaze-contingency that makes it particularly attractive for clinical and scientific applications (Thomas, Acton, Erichsen, Redmond, & Dunn, 2024).

Our method builds directly on these core improvements but enhances them substantially. It offers an order-of-magnitude increase in spatial and temporal precision, enabling reliable and repeatable measurements of visual function within the central 1° of the visual field. Even in the most advanced microperimetry systems, the entire foveola is typically represented by a single data point (Fig. 1E, Charng et al., 2020). It has even been argued that the term *microperimetry* is somewhat of a misnomer, since both probe sizes and the tested visual field size are often comparable to that of standard perimetry (Yang & Dunbar, 2021).

By contrast, our method resolves functional structure within the foveola, opening the door to more sensitive assessments of central vision. These capabilities make our technique an attractive tool for clinical evaluation. Its gaze-contingent nature allows precise testing even in individuals with unstable or eccentric fixation, such as those with macular degeneration or neurological disorders (Alexander, Macknik, & Martinez-Conde, 2018; Wolf, Ueda, & Hirano, 2021), and it enables direct quantification of fixational behavior as markers of disease severity or progression.

More broadly, the principles underlying our method could also improve perimetry testing at larger scales, addressing long-standing concerns about measurement noise in current protocols (Russell, Crabb, Malik, & Garway-Heath, 2012; Saunders, Russell, & Crabb, 2015). A major source of this noise stems from unaccounted eye movements, such as microsaccades, which can disrupt the alignment of stimuli and decrease test reliability (Pfau et al., 2021; Josan et al., 2023). By actively detecting and compensating for these movements, our system can achieve greater consistency across sessions (cf. Thomas et al., 2024), especially in individuals with high microsaccade rates (Figs. 6-7). Its potential extension to full-field assessments suggests broader applications across all types of visual function testing, delivering more accurate and individualized measures of sensitivity.

Beyond its clinical utility, our method provides a new tool for investigating fundamental questions in visual neuroscience. We observe that sensitivity patterns are highly idiosyncratic across individuals (Fig. 5A-B), yet remarkably stable over time (Fig. 7A). Contrary to dominant beliefs that visual sensitivity in the foveola is uniform and that peak sensitivity aligns with the center of gaze (Barten, 1999; Bedggood, Daaboul, Ashman, Smith, & Metha, 2008), we demonstrate that foveal sensitivity is highly non-uniform and that peak sensitivity deviates from the center of fixation. Since our observers manually calibrate their center of gaze to coincide with their subjective center of gaze, the center coordinate in our measures represent their preferred fixational locus. Interestingly, recent studies show that the preferred fixational locus is typically shifted 5 arcmin away from the topographical center of the retina towards the nasal visual field (Reiniger, Domdei, Holz, & Harmening, 2021). As a result, we might expect peak sensitivity in our measures to be shifted correspondingly. However, we found peak sensitivity to be shifted in the opposite direction and by much larger extents of 17 arcmin (Fig. 5C). These findings challenge existing models of central vision and suggest that anatomical factors, such as variability in the photoreceptor mosaic and photoreceptor density may contribute to the observed patterns (Li, Tiruveedhula, & Roorda, 2010; Hirsch & Curcio, 1989), though alternative explanations remain to be explored (Domdei, Reiniger, Holz, & Harmening, 2021).

Our system also enables detailed exploration on the role of fixational eye movements, including microsaccades and eye drift, in shaping visual function. We found that, irrespective of subject’s expertise, the presence of microsaccades was a significant factor in reducing visual sensitivity, a phenomenon likely linked to saccadic suppression (Intoy et al., 2021). This could have profound implications for understanding attention and perception (Poletti, 2023). Additionally, our method allows to precisely quantify eye drift, which has been linked to measures of visual acuity (Ratnam, Domdei, Harmening, & Roorda, 2017; Clark et al., 2022; Nghiem, Witten, Dufour, Harmening, & Azeredo da Silveira, 2025). This could pave the way for more nuanced assessments of visual function, providing new insights into how these small eye movements give rise to our rich visual percepts (Rucci & Victor, 2015).

In summary, we developed a method that enables precise, high-resolution mapping of visual function across the foveola. Our foveal perimetry combines cutting-edge eye tracking technology with real-time retinal stabilization, and rigorous psychophysical techniques. This innovative approach offers new insights into the spatial organization of foveal sensitivity and has significant implications for both basic visual neuroscience and clinical practice. By addressing the limitations of traditional perimetry methods, our system has the potential to become a new standard tool for evaluating visual function, advancing our understanding of vision across healthy and clinical populations alike.

## 4 Methods

### 4.1 Apparatus

We used an LCD monitor (ASUS PG259QN; 1920 x 1080 pixels; 543 x 303 mm) with a refresh rate of 360 Hz. Luminance was linearized across the entire output range of the monitor. Luminance measurements were performed with a Minolta CS-100 photometer (Konica Minolta, Tokyo, Japan). The monitor was placed at a distance 2200 mm from the subjects. To track eye movements, we used a digital camera (IO Industries Flare 12Mx180CX-NIR) with a pixel width of 5.5 µm and a sampling rate of 1000 Hz at 2 megapixels. The left eye of the subjects was patched. To eliminate head movements, subjects were placed on a bite bar and their heads were fixed. The software and experiment were implemented in Linux and C++. Data analysis was performed in Matlab R2022b and Python 3. We used the Palamedes toolbox 1.11.10 (Prins & Kingdom, 2018) for fitting psychometric functions. All code will be made available at the point of publication.

### 4.2 Subjects

Eleven subjects (6 female, aged 24-35 years) participated in the experiment. Five subjects had normal vision, three subjects had corrected-to-normal vision, and another three subjects were myopic (20/40 acuity) and uncorrected. Visual acuity was determined with a Snellen eye chart. We included subjects with different eyesight to test to which extent the proposed foveal perimetry can be performed with different observers and how much their eyesight influences the resulting sensitivity measurements. This research study was approved by the University of Rochester’s Research Subjects Review Board.

### 4.3 Data collection

We used the method of constant stimuli to fit full psychometric curves at each location and for each subject with 5 contrast levels which were chosen individually for each observer based on prior piloting. Different contrasts were used across observer, but within individuals we used the same 5 contrasts across retinal locations. Weber contrasts varied between 0.1-0.81 (*∼*5-41% monitor contrast).

Subjects participated in 2-8 experimental sessions, each lasting 1-3 hours. Each session started with a practice block (65 trials) to get familiar with the task and setup. Afterwards, subjects participated in multiple experimental blocks, lasting 7-8 minutes on average.

Since trial initiation was self-paced, there was considerable variability in the number of trials that a subject could do comfortably in one block. Based on their pace and preference, we therefore set the number of trials per block to 195 (13 locations x 5 contrasts x 3 repetitions) or 325 (5 repetitions). Regardless of the number of repetitions per block, we collected data for each subject until we reached 60 repetitions for each contrast tested (i.e. 3900 trials) after trial exclusion. Trials were excluded if blinks or microsaccades occurred within a 50 ms period of the probe interval. In addition, trials were excluded if gaze reached outside 1° (radius) of the display center to avoid any interference of the fixation arcs (2.5° radius). The amount of excluded trials ranged from 1-23% for different subjects, mainly because of inter-individual differences in microsaccade rates (Fig. 6C).

### 4.4 Calculating threshold estimates

For each subject, we fitted multiple psychometric functions to the psychophysical data using a maximum-likelihood criterion. Psychometric functions were logistic functions on log-contrasts. We first fitted one psychometric function using the pooled data of all retinal locations. We set the lower asymptote of this function to the false alarm rate (i.e. the percentage of incorrect responses in flash-absent trials), and estimated the mean threshold, the slope and the lapse rate of each subject. We then fitted psychometric functions for each retinal location separately using the same parameter values for the lower asymptote, the slope and lapse rate. Hence, differences between psychometric functions within one subject were merely caused by differences in local thresholds. We validated the quality of the fits by the means of goodness-of-fit statistics.

### 4.5 Reliability of measurements

To estimate reliability numerically, we computed (1) the split-half reliability of our dataset, and (2) the SEM (standard error of means) of our measurements using bootstrapping.

Split-half reliability is a standard metric to assess the internal consistency and reliability of an instrument. To compute the split-half reliability, we randomly split our dataset into two halves, computed all thresholds (for each retinal location and each subject) for each half separately and finally correlated the threshold estimates between the two halves. For this, we used the standard formula for split-half reliability (Spearman-Brown formula):

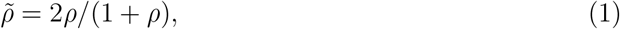

where 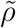 is the split-half reliability and *ρ* is the Pearson-correlation between thresholds. Split-half reliability was excellent (*ρ* = 0.97; Fig. 7B).

In addition, we computed the SEM of our threshold measurements. For this, we repeatedly (500 bootstraps) sampled different proportions of our data and computed all thresholds (for each location and each subject). We then computed how variable these threshold estimates are across bootstraps (=SEM) for each location and subject individually, and report their mean and variability.

## 5 Acknowledgments

This work was funded by NIH grant EY018363.

## Notes

### Competing Interest Statement

The authors have declared no competing interest.

## References

Acton, J. H., & Greenstein, V. C. (2013). Fundus-driven perimetry (microperimetry) compared to conventional static automated perimetry: similarities, differences, and clinical applications. Canadian Journal of Ophthalmology, 48 (5), 358–363. doi: 10.1016/j.jcjo.2013.03.021

Alexander, R. G., Macknik, S. L., & Martinez-Conde, S. (2018). Microsaccade characteristics in neurological and ophthalmic disease. Frontiers in Neurology, 9, 144. doi: 10.3389/fneur.2018.00144

Barten, P. G. (1999). Contrast sensitivity of the human eye and its effects on image quality. SPIE press. doi: 10.6100/IR523072

Bedggood, P., Daaboul, M., Ashman, R., Smith, G., & Metha, A. (2008). Characteristics of the human isoplanatic patch and implications for adaptive optics retinal imaging. Journal of Biomedical Optics, 13 (2), 024008–024008. doi: 10.1117/1.2907211

Bengtsson, B., & Heijl, A. (1998). SITA fast, a new rapid perimetric threshold test. Description of methods and evaluation in patients with manifest and suspect glaucoma. Acta Ophthalmologica Scandinavica, 76 (4), 431–437. doi: 10.1034/j.1600-0420.1998.760408.x

Bengtsson, B., Olsson, J., Heijl, A., & Rootzen, H. (1997). A new generation of algorithms for computerized threshold perimetry, SITA. Acta Ophthalmologica Scandinavica, 75 (4), 368–375. doi: 10.1111/j.1600-0420.1997.tb00392.x

Blackwell, H. R. (1952). Studies of psychophysical methods for measuring visual thresholds. JOSA, 42 (9), 606–616. doi: 10.1364/JOSA.42.000606

Bowers, N. R., Gautier, J., Lin, S., & Roorda, A. (2021). Fixational eye movements in passive versus active sustained fixation tasks. Journal of Vision, 21 (11), 16–16. doi: 10.1167/jov.21.11.16

Carr, J. W., Pescuma, V. N., Furlan, M., Ktori, M., & Crepaldi, D. (2022). Algorithms for the automated correction of vertical drift in eye-tracking data. Behavior Research Methods, 54 (1), 287–310. doi: 10.3758/s13428-021-01554-0

Cassels, N. K., Wild, J. M., Margrain, T. H., Chong, V., & Acton, J. H. (2018). The use of microperimetry in assessing visual function in age-related macular degeneration. Survey of Ophthalmology, 63 (1), 40–55. doi: 10.1016/j.survophthal.2017.05.007

Charng, J., Sanfilippo, P. G., Attia, M. S., Dolliver, M., Arunachalam, S., Chew, A. L., … Chen, F. K. (2020). Interpreting MAIA microperimetry using age-and retinal loci-specific reference thresholds. Translational Vision Science & Technology, 9 (7), 19–19. doi: 10.1167/tvst.9.7.19

Cherici, C., Kuang, X., Poletti, M., & Rucci, M. (2012). Precision of sustained fixation in trained and untrained observers. Journal of Vision, 12 (6), 31–31. doi: 10.1167/12.6.31

Choe, K. W., Blake, R., & Lee, S.-H. (2016). Pupil size dynamics during fixation impact the accuracy and precision of video-based gaze estimation. Vision Research, 118, 48–59. doi: 10.1016/j.visres.2014.12.018

Clark, A. M., Intoy, J., Rucci, M., & Poletti, M. (2022). Eye drift during fixation predicts visual acuity. Proceedings of the National Academy of Sciences, 119 (49), e2200256119. doi: 10.1073/pnas.2200256119

Clark, A. M., Moon, B., Jenks, S. K., Kapisthalam, S., & Poletti, M. (2023). The impact of sub-foveal scotomas on visual perception and fine oculomotor behavior. Journal of Vision, 23 (9), 5268–5268. doi: 10.1167/jov.23.9.5268

Cornsweet, T. N., & Crane, H. D. (1973). Accurate two-dimensional eye tracker using first and fourth Purkinje images. JOSA, 63 (8), 921–928. doi: 10.1364/JOSA.63.000921

Crane, H. D., & Steele, C. M. (1985). Generation-V dual-Purkinje-image eyetracker. Applied Optics, 24 (4), 527–537. doi: 10.1364/AO.24.000527

Daien, V., Peres, K., Villain, M., Colvez, A., Carriere, I., & Delcourt, C. (2014). Visual acuity thresholds associated with activity limitations in the elderly. The Pathologies Oculaires Liees a l’Age study. Acta Ophthalmologica, 92 (7), e500–e506. doi: 10.1111/aos.12335

Daniel, P. M., & Whitteridge, D. (1961). The representation of the visual field on the cerebral cortex in monkeys. The Journal of Physiology, 159 (2), 203. doi: 10.1113/jphysiol.1961.sp006803

Deubel, H., & Bridgeman, B. (1995). Fourth Purkinje image signals reveal eye-lens deviations and retinal image distortions during saccades. Vision Research, 35 (4), 529–538. doi: 10.1016/0042-6989(94)00146-D

Domdei, N., Reiniger, J. L., Holz, F. G., & Harmening, W. M. (2021). The relationship between visual sensitivity and eccentricity, cone density and outer segment length in the human foveola. Investigative Ophthalmology & Visual Science, 62 (9), 31–31. doi: 10.1167/iovs.62.9.31

Green, D. G. (1970). Regional variations in the visual acuity for interference fringes on the retina. The Journal of Physiology, 207 (2), 351. doi: 10.1113/jphysiol.1970.sp009065

Hawken, M. J., & Gegenfurtner, K. R. (2001). Pursuit eye movements to second-order motion targets. JOSA A, 18 (9), 2282–2296. doi: 10.1364/JOSAA.18.002282

Hazel, C. A., Petre, K. L., Armstrong, R. A., Benson, M. T., & Frost, N. A. (2000). Visual function and subjective quality of life compared in subjects with acquired macular disease. Investigative Ophthalmology & Visual Science, 41 (6), 1309–1315.

Heijl, A., Patella, V. M., & Bengtsson, B. (2021). Excellent perimetry - the field analyzer primer. Germany, Jena.

Hirsch, J., & Curcio, C. A. (1989). The spatial resolution capacity of human foveal retina. Vision Research, 29 (9), 1095–1101. doi: 10.1016/0042-6989(89)90058-8

Intoy, J., Mostofi, N., & Rucci, M. (2021). Fast and nonuniform dynamics of perisaccadic vision in the central fovea. Proceedings of the National Academy of Sciences, 118 (37), e2101259118. doi: 10.1073/pnas.2101259118

Johnson, C. A., Wall, M., & Thompson, H. S. (2011). A history of perimetry and visual field testing. Optometry and Vision Science, 88 (1), E8–E15. doi: 10.1097/OPX.0b013e3182004c3b

Josan, A. S., Farrance, I., Taylor, L. J., Adeyoju, D., Buckley, T. M., Jolly, J. K., & MacLaren, R. E. (2023). Microperimetry reliability assessed from fixation performance. Translational Vision Science & Technology, 12 (5), 21–21. doi: 10.1167/tvst.12.5.21

Kilpelainen, M., Putnam, N. M., Ratnam, K., & Roorda, A. (2021). The retinal and perceived locus of fixation in the human visual system. Journal of Vision, 21 (11), 9–9. doi: 10.1167/jov.21.11.9

Kimmel, D. L., Mammo, D., & Newsome, W. T. (2012). Tracking the eye non-invasively: simultaneous comparison of the scleral search coil and optical tracking techniques in the macaque monkey. Frontiers in Behavioral Neuroscience, 6, 49. doi: 10.3389/fnbeh.2012.00049

Laishram, M., Srikanth, K., Rajalakshmi, A. R., Nagarajan, S., & Ezhumalai, G. (2017). Microperimetry–a new tool for assessing retinal sensitivity in macular diseases. Journal of Clinical and Diagnostic Research: JCDR, 11 (7), NC08. doi: 10.7860/JCDR/2017/25799.10213

Lebedev, D. S., Belokopytov, A. V., Rozhkova, G. I., Vasilyeva, N. N., & Gracheva, M. A. (2024). Use of fixation microsaccades to increase the quality of visible images in the foveal zone. Neuroscience and Behavioral Physiology, 54 (9), 1488–1500. doi: 10.1007/s11055-024-01747-y

Legge, G. E., Rubin, G. S., Pelli, D. G., & Schleske, M. M. (1985). Psychophysics of reading — II. Low vision. Vision Research, 25 (2), 253–265. doi: 10.1016/0042-6989(85)90118-X

Li, K. Y., Tiruveedhula, P., & Roorda, A. (2010). Intersubject variability of foveal cone photoreceptor density in relation to eye length. Investigative Ophthalmology & Visual Science, 51 (12), 6858–6867. doi: 10.1167/iovs.10-5499

Nghiem, T. A., Witten, J. L., Dufour, O., Harmening, W. M., & Azeredo da Silveira, R. (2025). Fixational eye movements as active sensation for high visual acuity. Proceedings of the National Academy of Sciences, 122 (6), e2416266122. doi: 10.1073/pnas.2416266122

Pfau, M., Jolly, J. K., Wu, Z., Denniss, J., Lad, E. M., Guymer, R. H., … Schmitz-Valckenberg, S. (2021). Fundus-controlled perimetry (microperimetry): application as outcome measure in clinical trials. Progress in Retinal and Eye Research, 82, 100907. doi: 10.1016/j.preteyeres.2020.100907

Phu, J., Khuu, S. K., Yapp, M., Assaad, N., Hennessy, M. P., & Kalloniatis, M. (2017). The value of visual field testing in the era of advanced imaging: clinical and psychophysical perspectives. Clinical and Experimental Optometry, 100 (4), 313–332. doi: 10.1111/cxo.12551

Poletti, M. (2023). An eye for detail: Eye movements and attention at the foveal scale. Vision Research, 211, 108277. doi: 10.1016/j.visres.2023.108277

Poletti, M., Listorti, C., & Rucci, M. (2013). Microscopic eye movements compensate for nonhomogeneous vision within the fovea. Current Biology, 23 (17), 1691–1695. doi: 10.1016/j.cub.2013.07.007

Prins, N., & Kingdom, F. A. (2018). Applying the model-comparison approach to test specific research hypotheses in psychophysical research using the Palamedes toolbox. Frontiers in Psychology, 9, 1250. doi: 10.3389/fpsyg.2018.01250

Putnam, N. M., Hofer, H. J., Doble, N., Chen, L., Carroll, J., & Williams, D. R. (2005). The locus of fixation and the foveal cone mosaic. Journal of Vision, 5 (7), 3–3. doi: 10.1167/5.7.3

Qian, M., Wang, J., Gao, Y., Chen, M., Liu, Y., Zhou, D., … Roe, A. W. (2024). Multiple loci for foveolar vision in macaque monkey visual cortex. Nature Neuroscience, 1–13. doi: 10.1038/s41593-024-01810-4

Ratliff, F., & Riggs, L. A. (1950). Involuntary motions of the eye during monocular fixation. Journal of Experimental Psychology, 40 (6), 687. doi: 10.1037/h0057754

Ratnam, K., Domdei, N., Harmening, W. M., & Roorda, A. (2017). Benefits of retinal image motion at the limits of spatial vision. Journal of Vision, 17 (1), 30–30. doi: 10.1167/17.1.30

Reiniger, J. L., Domdei, N., Holz, F. G., & Harmening, W. M. (2021). Human gaze is systematically offset from the center of cone topography. Current Biology, 31 (18), 4188–4193. doi: 10.1016/j.cub.2021.07.005

Rentschler, I., & Treutwein, B. (1985). Loss of spatial phase relationships in extrafoveal vision. Nature, 313 (6000), 308–310. doi: 10.1038/313308a0

Rohrschneider, K., Bultmann, S., & Springer, C. (2008). Use of fundus perimetry (microperimetry) to quantify macular sensitivity. Progress in Retinal and Eye Research, 27 (5), 536–548. doi: 10.1016/j.preteyeres.2008.07.003

Rossi, E. A., & Roorda, A. (2010). The relationship between visual resolution and cone spacing in the human fovea. Nature Neuroscience, 13 (2), 156–157. doi: 10.1038/nn.2465

Rucci, M., & Victor, J. D. (2015). The unsteady eye: an information-processing stage, not a bug. Trends in Neurosciences, 38 (4), 195–206. doi: 10.1016/j.tins.2015.01.005

Russell, R. A., Crabb, D. P., Malik, R., & Garway-Heath, D. F. (2012). The relationship between variability and sensitivity in large-scale longitudinal visual field data. Investigative Ophthalmology & Visual Science, 53 (10), 5985–5990. doi: 10.1167/iovs.12-10428

Santini, F., Redner, G., Iovin, R., & Rucci, M. (2007). EyeRIS: a general-purpose system for eye-movement-contingent display control. Behavior Research Methods, 39 (3), 350–364. doi: 10.3758/BF03193003

Saunders, L. J., Russell, R. A., & Crabb, D. P. (2015). Measurement precision in a series of visual fields acquired by the standard and fast versions of the swedish interactive thresholding algorithm: analysis of large-scale data from clinics. JAMA Ophthalmology, 133 (1), 74–80. doi: 10.1001/jamaophthalmol.2014.4237

Sheehy, C. K., Yang, Q., Arathorn, D. W., Tiruveedhula, P., de Boer, J. F., & Roorda, A. (2012). High-speed, image-based eye tracking with a scanning laser ophthalmoscope. Biomedical Optics Express, 3 (10), 2611–2622. doi: 10.1364/BOE.3.002611

Shirato, S., Inoue, R., Fukushima, K., & Suzuki, Y. (1999). Clinical evaluation of SITA: a new family of perimetric testing strategies. Graefe’s Archive for Clinical and Experimental Ophthalmology, 237, 29–34. doi: 10.1007/s004170050190

SR Research. (2005). EyeLink II user manual, v2.14. Mississauga, ON.

SR Research. (2013). EyeLink 1000 plus user manual, v1.0.12. Mississauga, ON.

Stevenson, S. B., Roorda, A., & Kumar, G. (2010). Eye tracking with the adaptive optics scanning laser ophthalmoscope. In Proceedings of the 2010 symposium on eye-tracking research & applications (pp. 195–198). doi: 10.1145/1743666.1743714

Sunness, J. S., Gonzalez-Baron, J., Applegate, C. A., Bressler, N. M., Tian, Y., Hawkins, B., … Bergman, A. (1999). Enlargement of atrophy and visual acuity loss in the geographic atrophy form of age-related macular degeneration. Ophthalmology, 106 (9), 1768–1779. doi: 10.1016/S0161-6420(99)90340-8

Thomas, N., Acton, J. H., Erichsen, J. T., Redmond, T., & Dunn, M. J. (2024). Reliability of gaze-contingent perimetry. Behavior Research Methods, 56 (5), 4883–4892. doi: 10.3758/s13428-023-02225-y

Toet, A., & Levi, D. M. (1992). The two-dimensional shape of spatial interaction zones in the parafovea. Vision Research, 32 (7), 1349–1357. doi: 10.1016/0042-6989(92)90227-A

Tootell, R. B., Silverman, M. S., Switkes, E., & De Valois, R. L. (1982). Deoxyglucose analysis of retinotopic organization in primate striate cortex. Science, 218 (4575), 902–904. doi: 10.1126/science.7134981

VPixx Technologies. (2020). TRACKpixx3 installation guide, v1.4. Saint-Bruno, QC.

Vullings, C., & Verghese, P. (2021). Mapping the binocular scotoma in macular degeneration. Journal of Vision, 21 (3), 9–9. doi: 10.1167/jov.21.3.9

Wang, J. Z., Cherici, C., & Rucci, M. (2025). Spatial and temporal factors influencing fixational saccades. Journal of Neuroscience, 45 (39). doi: 10.1523/JNEUROSCI.2175-24.2025

Weymouth, F. W. (1958). Visual sensory units and the minimal angle of resolution. American Journal of Ophthalmology. doi: 10.1016/0002-9394(58)90042-4

Wolf, A., Ueda, K., & Hirano, Y. (2021). Recent updates of eye movement abnormalities in patients with schizophrenia: A scoping review. Psychiatry and Clinical Neurosciences, 75 (3), 82–100. doi: 10.1111/pcn.13188

Wu, R. J., Clark, A. M., Cox, M. A., Intoy, J., Jolly, P. C., Zhao, Z., & Rucci, M. (2023). High-resolution eye-tracking via digital imaging of Purkinje reflections. Journal of Vision, 23 (5), 4–4. doi: 10.1167/jov.23.5.4

Yang, Y., & Dunbar, H. (2021). Clinical perspectives and trends: microperimetry as a trial endpoint in retinal disease. Ophthalmologica, 244 (5), 418–450. doi: 10.1159/000515148

